# Arkas: Rapid, Reproducible RNAseq Analysis as a Service

**DOI:** 10.1101/031435

**Authors:** Anthony Colombo, Timothy J. Triche, Giridharan Ramsingh

**Affiliations:** Jane Anne Nohl Division of Hematology, Keck School of Medicine of USC, Los Angeles, California, USA

**Keywords:** transcription, sequencing, RNAseq, reproducibility, automation, cloud computing

## Abstract

The recently introduced Kallisto[1] pseudoaligner has radically simplified the quantification of transcripts in RNA-sequencing experiments. However, as with all computational advances, reproducibility across experiments requires attention to detail. The elegant approach of Kallisto reduces dependencies, but we noted differences in quantification between versions of Kallisto, and both upstream preparation and downstream interpretation benefit from an environment that enforces a requirement for equivalent processing when comparing groups of samples. Therefore, we created the Arkas[3] and TxDbLite[4] R packages to meet these needs and to ease cloud-scale deployment of the above. TxDbLite extracts structured information directly from source FASTA files with per-contig metadata, while Arkas enforces versioning of the derived indices and annotations, to ensure tight coupling of inputs and outputs while minimizing external dependencies. The two packages are combined in Illumina's BaseSpace cloud computing environment to offer a massively parallel and distributed quantification step for power users, loosely coupled to biologically informative downstream analyses via gene set analysis (with special focus on Reactome annotations for ENSEMBL transcriptomes). Previous work (e.g. Soneson et al., 2016[34]) has revealed that filtering transcriptomes to exclude lowly-expressed isoforms can improve statistical power, while more-complete transcriptome assemblies improve sensitivity in detecting differential transcript usage. Based on earlier work by Bourgon et al., 2010[11], we included this type of filtering for both gene- and transcript-level analyses within Arkas. For reproducible and versioned downstream analysis of results, we focused our efforts on ENSEMBL and Reactome[2] integration within the qusage[19] framework, adapted to take advantage of the parallel and distributed environment in Illumina’s BaseSpace cloud platform. We show that quantification and interpretation of repetitive sequence element transcription is eased in both basic and clinical studies by just-in-time annotation and visualization. The option to retain pseudoBAM output for structural variant detection and annotation, while not insignificant in its demand for computation and storage, nonetheless provides a middle ground between *de novo* transcriptome assembly and routine quantification, while consuming a fraction of the resources used by popular fusion detection pipelines and providing options to quantify gene fusions with known breakpoints without reassembly. Finally, we describe common use cases where investigators are better served by cloud-based computing platforms such as BaseSpace due to inherent efficiencies of scale and enlightened common self-interest. Our experiences suggest a common reference point for methods development, evaluation, and experimental interpretation.

## 1 Introduction

The scale and complexity of sequencing data in molecular biology has exploded in the 15 years following completion of the Human Genome Project[36]. Furthermore, as a dizzying array of *-Seq protocols have been developed to open new avenues of investigation, a much broader cross-section of biologists, physicians, and computer scientists have come to work with biological sequence data. The nature of gene regulation (or, perhaps more appropriately, transcription regulation), along with its relevance to development and disease, has undergone massive shifts propelled by novel approaches, such as the discovery of evolutionarily conserved non-coding RNA by enrichment analysis of DNA[35] and isoform-dependent switching of protein interactions. Sometimes lost within this excitement, however, is the reality that biological interpretation these results can be highly dependent upon both their extraction and annotation. A rapid, memory-efficient approach to estimate abundance of both known and putative transcripts substantially broadens the scope of experiments feasible for a non-specialized laboratory and of methodological work deemed worthwhile given the pace of innovations. Recent work on the Kallisto pseudoaligner, among other k-mer based approaches, has resulted in exactly such an approach.

In order to leverage this advance for our own needs, which have included the quantification of repetitive element transcription in clinical trials, the comparison of hundreds of pediatric malignancies with their adult counterparts, and the analysis of single-cell RNA sequencing benchmarks, we first created an R package (Arkas) which automates the construction of composite transcriptomes from multiple sources, along with their digestion into a lightweight analysis environment. Further work led us to automate the extraction of genomic and functional annotations directly from FASTA contig comments, eliding sometimes-unreliable dependencies on services such as Biomart. Finally, in a nod to enlightened self-interest and mutual benefit, we collaborated with Illumina to deploy the resulting pipeline within their BaseSpace cloud computing platform, largely to avoid internal bureaucratic inertia and increase uptake of the resulting tools. The underlying packages are freely available with extensive use-case vignettes on GitHub, are in preparation for Bioconductor submission and review, and available in beta for review upon request, pending full public BaseSpace deployment.

High-performance computing (HPC) based bioinformatic workflows have three main subfamilies: in-house computational packages, virtual-machines, and cloud based computational environments. The in-house approaches are substantially less expensive when raw hardware is in constant use and dedicated support is available, but internal dependencies can limit reproducibility of computational experiments. Specifically, superuser access needed to deploy container-based, succinct code encapsulations (often referred to as “microservices” elsewhere) can run afoul of normal permissions, and the maintenance of a broadly usable set of libraries across nodes for diverse users can lead to shared object code linking dynamically to different libraries under different user environments. Virtual machines (VMs) have similar issues related to maintenance, as a heavyweight machine image is required for each instance, though the consequences of privelege escalation are decreased.

By contrast, modern cloud-based approaches to distributed and parallel computing are forced by necessity to offer a lowest-common-denominator platform with high availability to the broadest possible user base. Platform-as-a-service approaches take this one step further, offering controlled deployment and fault tolerance across potentially unreliable instances provided by third parties (Amazon Elastic Compute Cloud(EC2), in the case of BaseSpace) and enforcing a standard for encapsulation of developers’ services (Docker, in the case of BaseSpace). Within this framework, the user or developer cedes some control of the platform and interface, in exchange for the platform provider handling the details of workflow distribution and execution. In our recent experience, this has provided the best compromise of usability and reproducibility when dealing with users of widely varying skill levels, and the lightweight-container approach exemplified by Docker leads to a more rapid pace of development and deployment than is possible with VMs. Combined with versioning of deployments, it is feasible for a user to reconstruct results from an earlier point in time, while simultaneously re-evaluating the generated data under state-of-the-art implementations.

Several recent high impact publications have used cloud-computing work flows such as CloudBio-linux, CloudMap, and Mercury over AWS EC2 [27]. CloudBio-linux software is centered around comparative genomics, phylogenomics, transcriptomics, proteomics, and evolutionary genomics studies using Perl scripts [27]. Although offered with limited scalability, the CloudMap software allows scientists to detect genetic variations over a field of virtual machines operating in parallel [27]. For comparative genomic analysis, the Mercury [28] workflow can be deployed within Amazon EC2 through instantiated virtual machines but is limited to BWA and produces a variant call file (VCF) without considerations of pathway analysis or comparative gene set enrichment analyses. The effectiveness for conducting genomic research is greatly influenced by the choice of computational environment.

The majority of RNA-Seq analysis pipelines consist of read preparation steps, followed by computationally expensive alignment against a reference genome (which itself is not representative of the generating spliced sequences in the source analyte). Software for calculating transcript abundance and assembly can surpass 30 hours of computational time [1]. When known or putative transcripts of defined sequence are the primary interest, Kallisto[1] and colleagues have shown that near-optimal RNAseq transcript quantification is achievable in minutes minutes on a standard laptop. After verifying these numbers on our own laptops, we became interested in a massively parallel yet easy-to-use approach that would allow us to perform the same task on arbitrary datasets, and reliably interpret the output from colleagues doing the same. In collaboration with Illumina, we found that the freely available BaseSpace platform was already well suited for this purpose, with automated ingestion of SRA datasets as well as newly produced data from core facilities using recent Illumina sequencers. The design of our framework emphasizes loose coupling of components and tight coupling of reference transcriptome annotations; nonetheless, we find that the ease of use and massive parallelization provided by BaseSpace has made it our default execution environment.

We encapsulated the resulting packages, along with Kallisto itself, in Docker containers that perform transcript abundance quantification; a loosely coupled second step of the Arkas work-flow implements rapid set enrichment analysis over Illumina’s BaseSpace Platform.

The BaseSpace Platform utilizes AWS cc2 8x-large instances by default, each with access to eight 64-bit CPU cores and virtual storage of over 3 terabytes. Published applications on BaseSpace can allocate up to 100 such nodes, distributing analyses such that many samples can be processed simultaneously, in parallel. Direct imports of existing experiments from SRA, along with default availability of experimenters’ own reads, fosters a critical environment for independent replication and reanalysis of published data.

A second bottleneck in bioinformatic workflows, hinted at above, arises from the frequent transfer and copying of source data across local networks and/or the internet. With a standardized deployment platform, it becomes easier to move executable code to the environment of the target data, rather than transferring massive datasets into the environment where the executable workflows were developed. For example, an experiment from SRA with reads totaling 141.3GB is reduced to summary quantifications totaling 1.63GB (nearly two orders of magnitude) and a report of less than 10MB (a further two orders of magnitude), for a total reduction in size exceeding 4 orders of magnitude with little or no loss of user-visible information. Moreover, the untouched original data is never discarded unless the user explicitly demands it, something which can rarely be said of local compute environments, and the location of original sources is always traceable, again in contrast to local HPC.

## 2 Materials

### 2.1 Kallisto

Kallisto [1] quantifies transcript abundance from input RNA-Seq reads by using a process known as pseudoalignment which identifies the read-transcript compatibility matrix. The compatibility matrix is formed by counting the number of reads with the matching alignment; the equivalence class matrix has a much smaller dimension compared to matrices formed by transcripts and read coverage. Computational speed is gained by performing the Expectation Maximization [31] (EM) algorithm over a smaller matrix.

### 2.2 Arkas

Arkas[3] is a BioConductor package that automates the index caching, annotation, and quantification associated with running the Kallisto pseudoaligner integrated within the BioConductor environment.

Arkas can process raw reads into transcript- and pathway-level results within BioConductor or in Illuminas BaseSpace cloud platform.

Arkas[3] reduces the computational steps required to quantify and annotate large numbers of samples against large catalogs of transcriptomes. Arkas calls Kallisto[1] for on-the-fly transcriptome indexing and quantification recursively for numerous sample directories. For RNA-Seq projects with many sequenced samples, Arkas encapsulates expensive transcript quantification preparatory routines, while uniformly preparing Kallisto [1] execution commands within a versionized environment encouraging reproducible protocols.

Arkas [3] integrates quality control analysis for experiments that include Ambion spike-in controls defined from the External RNA Control Consortium [30][29]. Arkas includes erccdashboard [15], a BioConductor package for analyzing data sequenced with ERCC (Ambion) spike-ins. We designed a function titled ‘erccAnalysis’ which produces Receiver Operator Characteristic (ROC) Curves by preparing differential expression testing results for spike-in analysis.

Arkas imports the data structure from SummarizedExperiment[16] and creates a sub-class titled KallistoExperiment which preserves the S4 structure and is convenient for handling assays, phenotypic and genomic data. KallistoExperiment includes GenomicRanges[32], preserving the ability to handle genomic annotations and alignments, supporting efficient methods for analyzing high-throughput sequencing data. The KallistoExperiment sub-class serves as a general purpose container for storing feature genomic intervals and pseudoalignment quantification results against a reference genome called by Kallisto [1]. By default KallistoExperiment couples assay data such as the estimated counts, effective length, estimated median absolute deviation, and transcript per million count where each assay data is generated by a Kallisto [1] run; the stored feature data is a GRanges object from GenomicRanges[32], storing transcript length, GC content, and genomic intervals. Arkas[3] is a portable work-flow that includes a routine ‘SEtoKE‘ which casts a SummarizedExperiment [16] object into a KallistoExperiment object handy for general pathway analysis, transcript- and/or gene-wise analysis.

Given a KallistoExperiment, downstream enrichment analysis of bundle-aggregated transcript abundance estimates are performed using QuSage [19] imported from BioConductor. For gene-set enrichment analysis, Qusage [19] calculates the variance inflation factor which corrects the inter-gene correlation that results in high Type 1 errors using pooled, or non-pooled variances between sample groups. We’ve customized the Qusage [19] algorithm accelerating its computational speed improving its performance on average by 1.34X (Figure 5)(Table 6) 5. We improved QuSage’s [19] performance using RcppArmadillo[20] modifying only the calculations for the statistics defined for Welch’s test for unequal variances between sample groups; the shrinkage of the pooled variances is performed using the CAMERA algorithm within limma[17].

**Figure 5:**
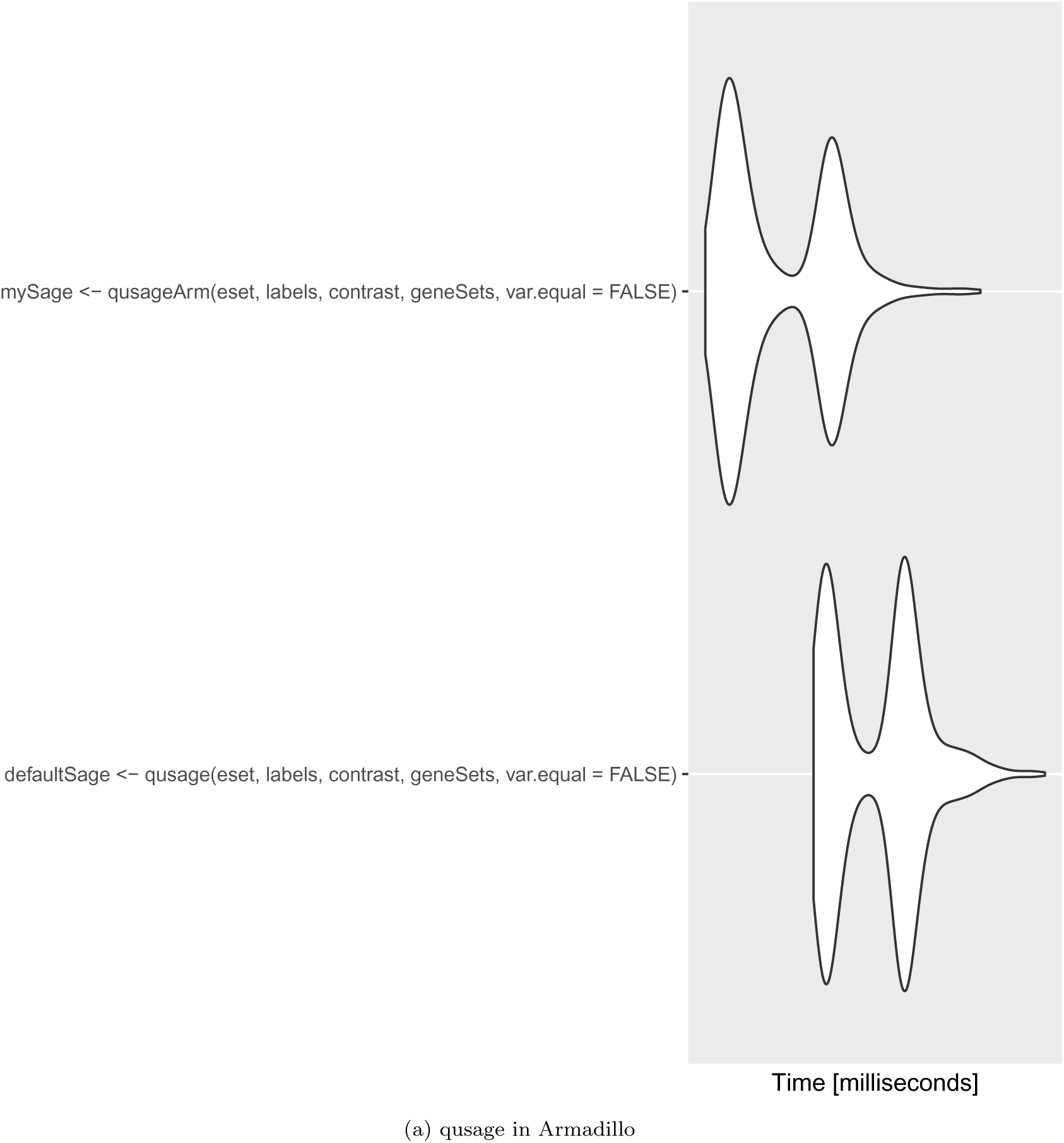
computational microbenchmark

**Table 1:** Variation of v.0.42.3

| Standard Deviation of percent differences between v.0.42.3 |  |  |  |
| --- | --- | --- | --- |
| Kallisto v.0.42.3 | Estimated Counts | Effective Length | Estimated MAD |
| Run1 | 0 | 0 | 0 |
| Run2 | 0 | 0 | 0 |
| Run3 | 0 | 0 | 0 |
| Run4 | 0 | 0 | 0 |
| Run5 | 0 | 0 | 0 |
| Run6 | 0 | 0 | 0 |
| Run7 | 0 | 0 | 0 |
| Run8 | 0 | 0 | 0 |
| Run9 | 0 | 0 | 0 |
| Run10 | 0 | 0 | 0 |

**Table 2:** Standard Deviation of Percent Error 0.42.1 and 0.42.3

| Standard Deviation of percent differences between v.0.42.1 and v.0.42.3 |  |  |  |  |
| --- | --- | --- | --- | --- |
| Kallisto v.0.42.3 and v.0.42.1 | Estimated Counts | Effective Length | Estimated MAD | TPM |
| Run1 | 419.7398 | 23.49247 | 74.54197 | 423.5493 |

**Table 3:** Arkas Run Time KallistoExperiment

| System Runtime for creating KallistoExperiment (secs) |  |  |
| --- | --- | --- |
| user | system | elapsed |
| 23.551 | 1.032 | 19.107 |

**Table 4:** Automated Annotation Run Time

| System Runtime for full Annotation of a merged KallistoExperiment (secs) |  |  |
| --- | --- | --- |
| user | system | elapsed |
| 2.336 | 0.064 | 2.397 |

**Table 5:**
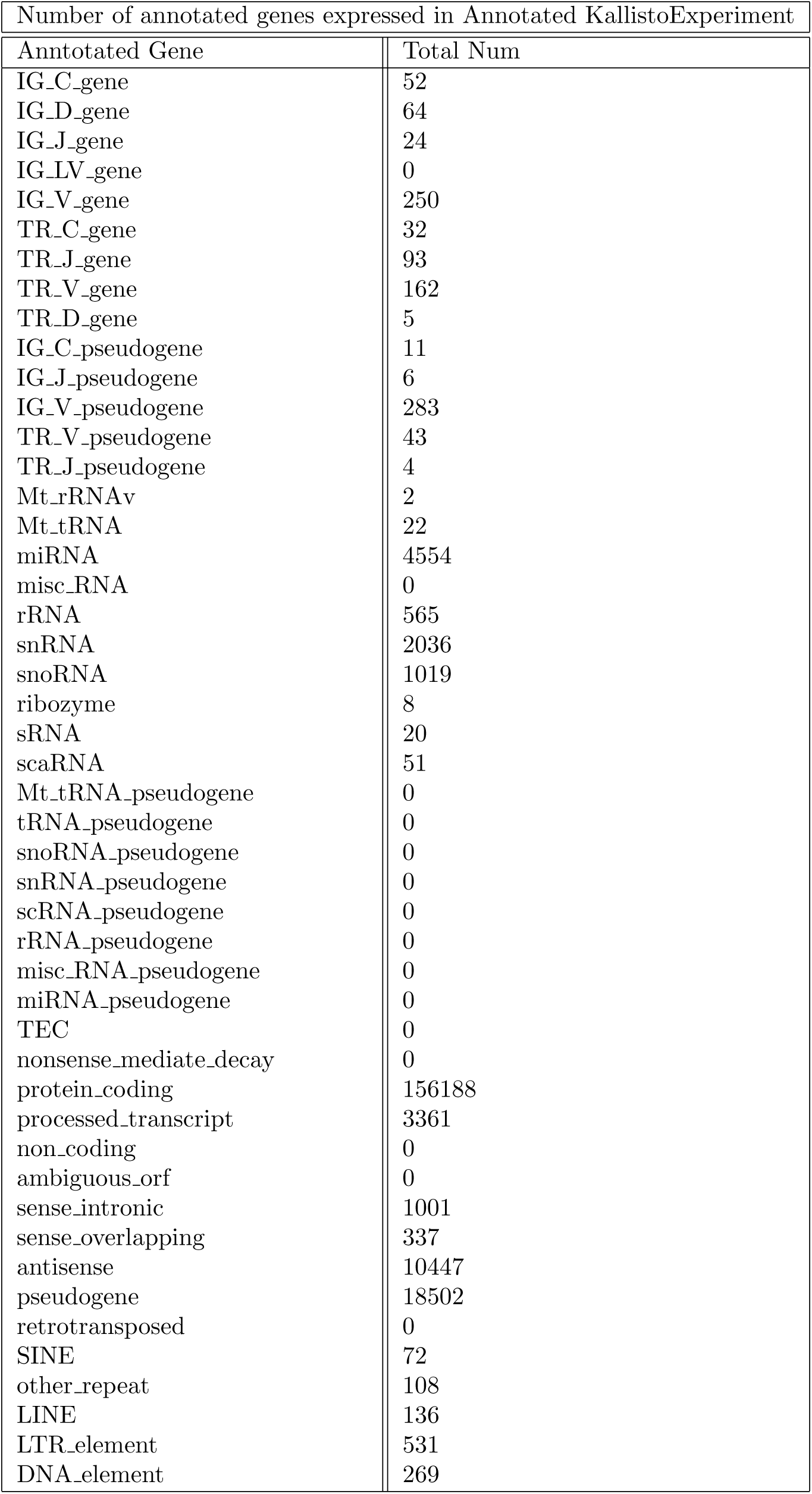
Summary of Genes Annotated from an Experiment

**Table 6:**
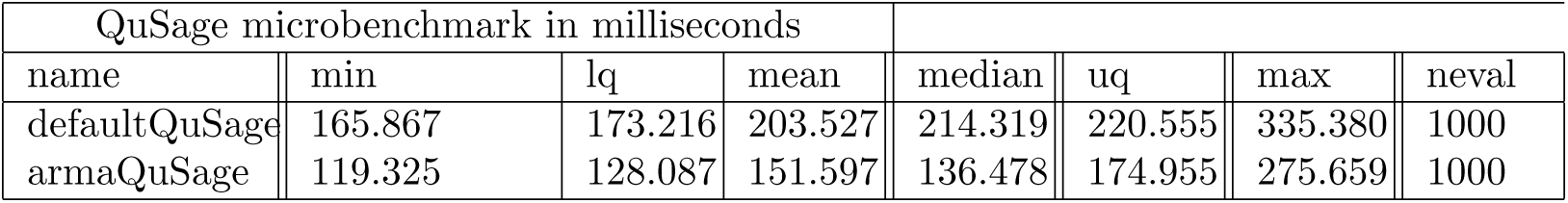
Qusage and Armadillo Benchmarks

Arkas’[3] accelerated enrichment analysis is useful for analyzing large gene-sets from Molecular Signatures DataBase MSigDB [21] Reactome [9], and/or other signature gene sets simultaneously.

Pathway enrichment analysis can be performed from the BaseSpace cloud system downstream from parallel differential expression analysis. BaseSpace Cloud Platform offers Advaita among one of the many published computational software publicly available. Advaita offers an extensive pathway analysis software available within the BaseSpace environment, so we’ve customized Arkas’ routine ‘geneWiseAnalysis.R’ to output differential expression formatted for downstream importing into Advaita.

### 2.3 TxDbLite

The choice of catalog, the type of quantification performed, and the methods used to assess differences can profoundly influence the results of sequencing analysis. Ensembl reference genomes are provided to GENCODE as a merged database from Havana’s manually curated annotations with Ensembl’s automatic curated coordinates. AceView, UCSC, RefSeq, and GENCODE have approximately twenty thousand protein coding genes, however AceView and GENCODE have a greater number of protein coding transcripts in their databases. RefSeq and UCSC references have less than 60,000 protein coding transcripts, whereas GENCODE has 140,066 protein coding loci. AceView has 160,000 protein coding transcripts, but this database is not manually curated [6]. The database selected for protein coding transcripts can influence the amount of annotation information returned when querying gene/transcript level databases.

Although previously overlooked, non-coding RNAs have been shown to share features and alternate splice variants with mRNA revealing that non-coding RNAs play a central role in metastasis, cell growth and cell invasion [24]. Non-coding transcripts have been shown to be functional and are associated with cancer prognosis; proving the importance of studying ncRNA transcripts.

Each non-coding database is curated at different frequencies with varying amounts of non-coding RNA enteries that influences that mapping rate. GENCODE Non-coding loci annotations contain 9640 loci, UCSC with 6056 and 4888 in RefSeq [6]; GENCODE annotations have the greatest number of lncRNA, protein and non-coding transcripts, and highest average transcripts per gene, with 91043 transcripts unique to GENCODE absent for UCSC and RefSeq databases [6]. Ensembl and AceView annotate more genes in comparison to RefSeq and UCSC, and return higher gene and isoform expression labeling improving differential expression analyses [7]. Ensembl achieves conspicuously higher mapping rates than RefSeq, and has been shown to annotate larger portions of specific genes and transcripts that RefSeq leaves unannotated [7]. Ensembl/GENCODE annotations are manually curated and updated more frequently than AceView. Further Ensembl has been shown to detect differentially expressed genes that is approximately equivalent to AceView [7].

Although Repetitive elements comprise over two-thirds of the human genome, repeat elements were considered non-functional and were not studied closely. Alu (Arthrobacter luteus) elements are a subfamily of repetitive elements that are roughly 300 base pairs long making up 11 % of the human genome [25]. Alu elements are the most abundant retro-transposable element, shown to increase genomic diversity impacting the evolution of primates and contributing to human genetic diseases [25]. Alu elements have been shown to influence genomic diversity through genetic instability. Regarding breast and ovarian tumorigenesis, the infamous BRCA1 and BRCA2 genes associated with survivability and prognosis contain genomic regions with very high densities of repetitive DNA elements [26].

Our understanding of biology is deepened when investigating expression of transcripts that include a merged transcriptome of coding, non-coding, and repetitive elements from data bases frequently curated. A *complete* transcriptome achieves higher mapping rates over larger catalogs of transcript families. We present TxDbLite[4] a fast genomic ranges annotation package that is included within the Arkas workflow that annotates non-coding, and repetitive elements on-the-fly downstream of transcript quantification steps.

TxDbLite [4] is a minimalist annotation package generator, designed to extract annotations from FASTA files. The underlying assumption is that users want to quantify transcript-level RNA expression, may or may not have a GTF file for each transcriptome, and would like to extract as much information as possible about the transcripts from what they do have. This in turn allows the Arkas[3] package to automatically determine what transcriptomes a dataset came from, whether those are already known to the package, and how to generate certain types of annotation for specific transcriptomes from Ensembl, RepBase, and ERCC (spike-in controls from Ambion).

## 3 Methods

### 3.1 Arkas

Illumina sequencers generate demultiplexed FASTQ files; Arkas [3] programmatically orders file inputs as required by Kallisto into sets of demultiplexed reads. The routine ‘runKallisto’[3] pairs the corresponding demultiplexed reads, builds and caches an index consisting of External RNA Control Consortium (ERCC), non-coding RNA, and coding RNA, & RepBase repeatome transcripts. Quantification is issued against the transcriptome per-sample abundance, and individual quantified samples are merged into a single KallistoExperiment from the ‘mergeKallisto’[3] routine. The merged KallistoExperiment identifies genes from ‘collapseBundles’[3] method which collapses the merged bundles of transcripts by the respective gene ID contained in the FASTA reference, and discards any transcript that had less than one count across all samples. The total count for a successfully collapsed transcript bundle is aggregated from individual transcripts within bundles that passed the filtering criteria. The collapsed bundles are annotated using routines ‘annotateFeatures’[3].

Standard approaches to modeling transcript-level differences in abundance rely upon having substantial numbers of replicates per condition. One of the novel features of Kallisto [1], implemented by Harold Pimentel (reference), is fast bootstrap sampling at the transcript level within the expectation-maximization algorithm. Arkas implements hooks to quantify the impact of this uncertainty in repeat elements and spike-in controls, where compositional analysis tools in the R environment [10] are pressed into service. Computational analysis with additive log-contrast formulations has long been standard in geology and other fields. We are exploring its use as a within-bundle method to quantify the most prominent isoform-centric impacts in an expression analysis.

After obtaining bundled transcript aggregated counts labeled by the any arbitrary FASTA reference, a Gene-wise analysis was performed. The gene level analysis is invoked from ‘geneWiseAnalysis’[3] which imports limma[17]) and edgeR [18] to normalize library sizes fitting an arbitrary marginal contrast, then propagates the resulting signed log10(p) values through clustered and un-clustered pathway enrichment analyses.

As previously, if the user has provided a grouping factor or design matrix, marginal significance for individual pathways and overall perturbation is assessed. Arkas discards transcripts and/or bundles with few reads (default 1) to improve statistical power [11].

### 3.2 BaseSpace

Arkas is currently written as a virtualized operating system, which can run on the BaseSpace platform generating the Kallisto pseudoaligned files. Arkas-BaseSpace can import SRA files and quantify transcript abundance.

### 3.3 TxDbLite

Gene annotation is performed from user-selected bundled transcriptomes (ERCC, Ensembl, and/or RepBase) simultaneously merging annotated samples into one R object: KallistoExperiment. We currently support reference databases for Homo sapiens and Mouse (NCBI). Routines such as ‘annotateBundles’ annotate transcriptomes from databases for example External RNA Control Consortium (ERCC), non-coding RNA, coding RNA, & RepBase repeatomes.

The design structure of Arkas versionizes the Kallisto [1] reference index to enforce that the Kallisto software versions are identical amidst merged KallistoExperiment data containers prior downstream analysis. Enforcing reference versions and Kallisto [1] versions prevents errors when comparing experiments. When the KallistoExperiment is generated, the Kallisto [1] version is stored within a data slot and can be accessed using the command ‘kallistoVersion(KallistoExperiment)’. Before kallisto quantified data can be merged, Arkas first checks the Kallisto [1] index name and version from the *runJnfo.json* file and enforces matching version.

### 3.4 Quality Control Using ERCC-SpikeIns (Ambion)

Arkas workflow integrates the BioConductor package ‘erccdashboard’ [15] which tests for quality, and false positive rates. If the library preparation includes ERCC-Spike Ins, Arkas will generate useful receiver operator characteristic plots, average ERCC Spike Amount volume, comparison plots of ERCC volume Amount and normalized ERCC counts. Arkas method ‘erccAnalysis’ also includes Normalized Spike In Amount against Percent Differences, and most significantly ‘erccAnalysis’ plots FPR vs. TPR (Figures 2, 3, 4) 2, 3 4

**Figure 2:**
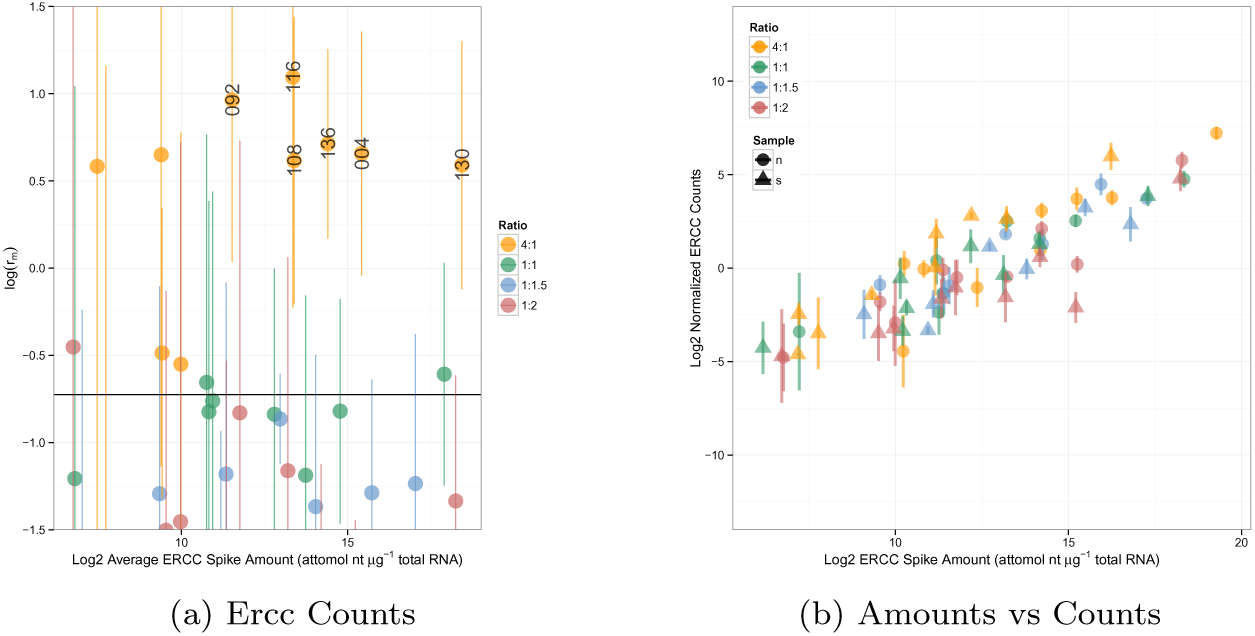
Ercc Counts

**Figure 3:**
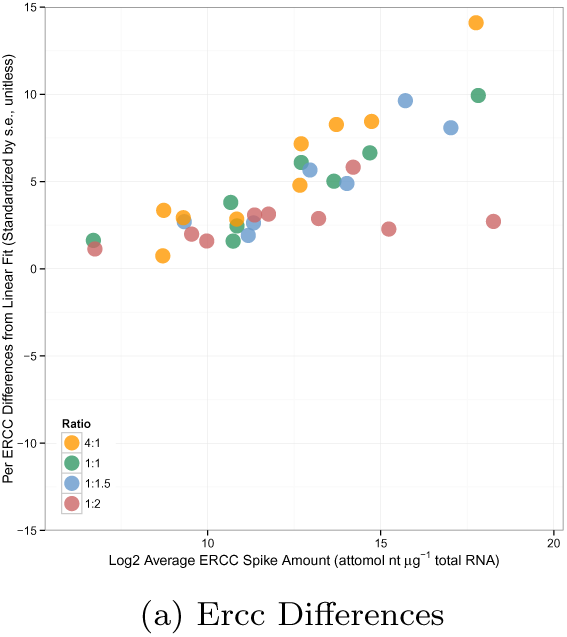
Ercc Differences

**Figure 4:**
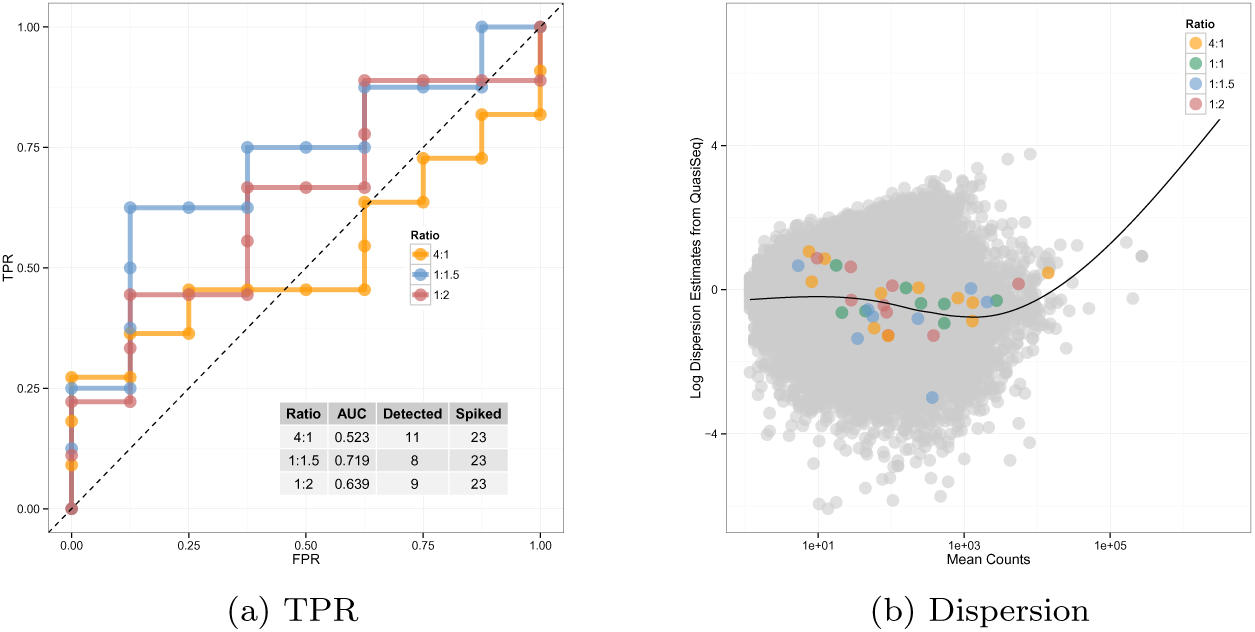
FPR vs TPR, and Dispersion

## 4 Results

### 4.1 Data Variance

Arkas enforces matching Kallisto versions in order run and merge Kallisto quantified data [1]. In order to show the importance of enforcing the same Kallisto version, we’ve repeatedly ran Kallisto on the same 6 samples, quantifying transcripts against two different Kallisto versions and measured the percent differences and standard deviation between these runs [1]. We ran Kallisto quantification once with Kallisto version 0.42.1, and 10 times with version 0.42.3 merging each run into a KallistoExperiment and storing the 11 runs into a list of Kallisto experiments [1].

We then analyzed the percent difference for each gene across all samples and calculated the standard deviation of version 0.42.3 of the 10 Kallisto runs generated by Arkas [1]. We randomly selected a KallistoExperiment v.0.42.3 from our KallistoExperiment list, and calculated the percent difference between each of the other 9 KallistoExperiments of the same version across all samples. The table 1 (Table 1) shows the standard deviation of the percent differences of the raw values such as estimated counts, effective length, and estimated median absolute deviation. Kallisto data quantified against the reference generated by the same kallisto version is 0 within the same version 0.42.3 for every transcript across all samples.

However, we compared the merged kallisto data of estimate raw abundance counts, effective length, estimated median absolute deviation, and transcript per million values between version 0.42.1 and a randomly selected KallistoExperiment data container generated by kallisto version 0.42.3. Table 2 (Table 2) shows that there exists large standard deviations of the percent differences calculated between each gene expression across all samples. This shows the importance of enforcing uniform versions.

We plotted the errors between Kallisto versions, and fit each of the calculated percent differences to a normally distributed data set generated by the mean and standard deviation of the percent differences of each assay data [1]. The QQ plots show that the errors are some-what normal (Figure 1); thus we confidently create default settings which prevent the public from analyzing or sharing data from different versions.

**Figure 1:**
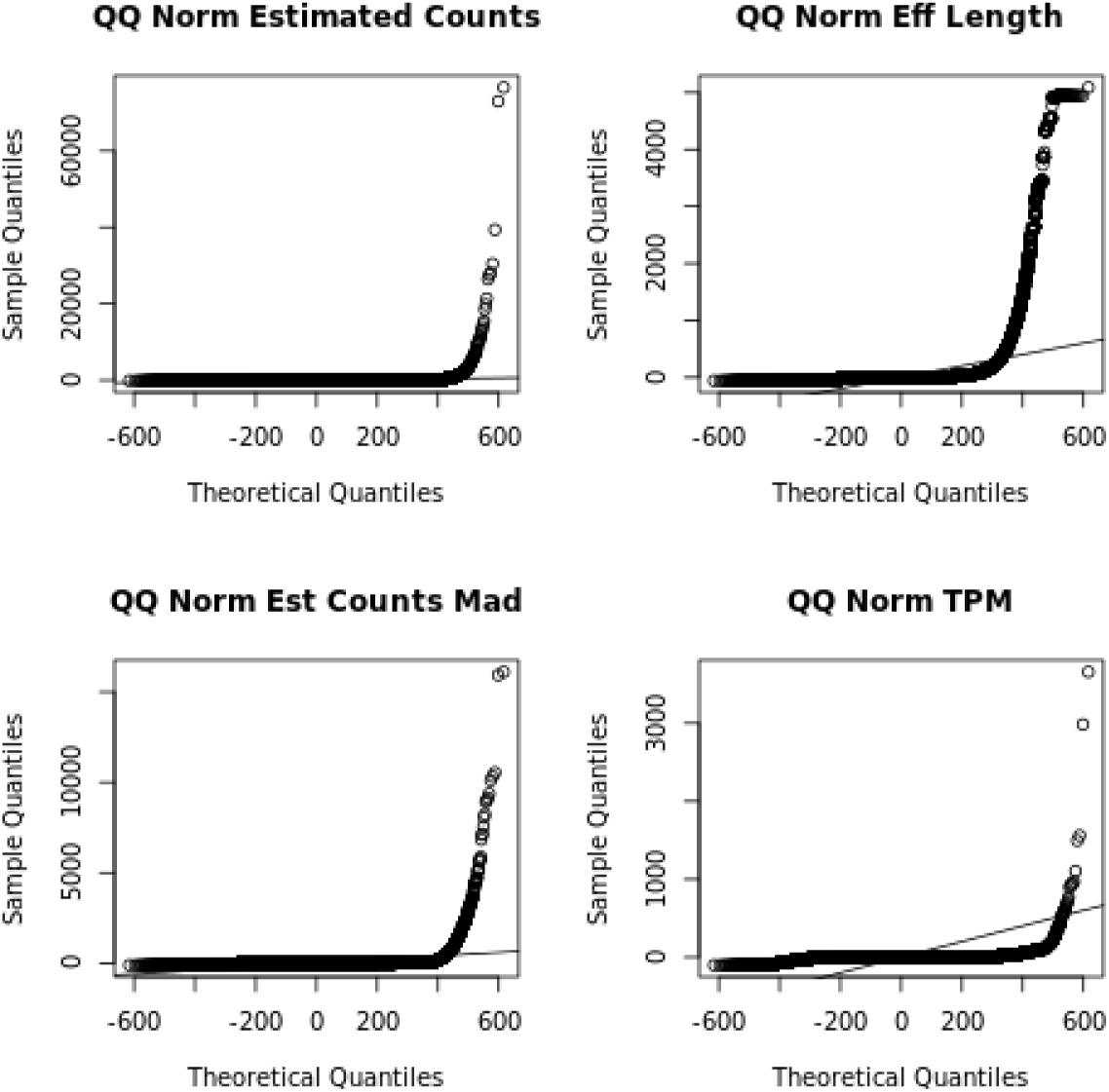
Quantile Plots of Percent Difference between v.0.42.3 and v.0.42.1

### 4.2 Annotation of Coding, non-Coding, and Repeat Elements

The annotations were performed with a run time of 2.336 seconds (Table 4) on a merged Kallisto [1] sample directory creating a KallistoExperiment class with feature data containing a GenomicRanges [32] object with 213782 ranges and 9 metadata columns. After the sample directories container raw fastq files were quantified using Arkas, the Arkas routine ‘mergeKallisto’ was performed against 6 quantified samples. The system runtime for creating a merged KallistoExperiment class for 6 samples was 23.551 seconds (Table 3).

### 4.3 ERCC-Analysis

The results for ERCC analysis includes 5 plots generated by Arkas’ method ‘erccAnalysis’, and the plots are saved calling ‘saveArkasPlots’. The Ercc plots figure 2, 3 4 titled ERCC Counts, Ercc Differences, and TPR vs FPR and Dispersion plots (Figures 2,3,4).

### 4.4 QuSage Acceleration Analysis

For testing a competitive revision of QuSage [19], we used the dataset from a study by Huang Y et al. (GEOID: GSE30550) which examines the expression of 17 patients before and 77 hours after exposure to the in uenza virus. This expression set was selected because it was described in the QuSage vignette [19]; so we selected the matching enrichment gene set and expression set used in QuSage BioConductor vignette [19], and developed accelerated computational performance while ensuring accuracy.

The calculations for Welch’s Approximation, such as standard deviation and degrees of freedom, where performed by Armadillo C++ libraries seamlessly integrated into the R environment using RcppArmadillo [20]. The gained computational speed was achieved from altering the following QuSage functions ‘calcVIF.R’, ‘makeComparisons.R’, and ‘calculateIndividualExpressions.R’ [19]. We ensured that each of the C++ scripts had at most machine error precision between QuSage [19] defaults and the altered RcppArmadillo [20] libraries.

## 5 Discussion

Many research publications include a written methodology section with varying degrees of detail and an included supplemental section; this is done as a symbol for attempting validation. Reproducible research is defined as a link between the global research community, defined as the set of all members and associated writings therein, to unique members of the global data environment, the set of all archived, published, clinical, sequencing and other experimental data. The aim for practicing transparent research methodologies is to clearly define associations with every research experiment minimizing opaqueness between the analytical findings, clinical studies, and utilized methods. As for clinical studies, re-generating an experimental environment has a very low success rate [33] where non-validated preclinical experiments spawned developments of best practices for critical experiments. Re-creating a clinical study has many challenges including an experimental design that has a broad focus applicability, the difficult nature of a disease, complexity of cell-line models between mouse and human that creates an inability to capture human tumor environment, and limited power through small enrollments during the patient selection process [33]. Confirming preclinical data is difficult, however the class of re-validated experiments each contain carefully selected reagents, diligently tested controls for various levels of an experiment, and, most significantly, a complete description of the entire data set. Original data sets are frequently not reported in the final paper and usually removed during the revision process. Experimental validation is dependent on the skillful performance of an experiment, and an earnest distribution of the analytic methodology which should contain most, if not all, raw and resultant data sets.

With recent developments for virtualized operating systems, developing best practices for bioinformatic confirmations of experimental methodologies is much more straightforward in contrast to duplicating clinical trials’ experimental data for drug-development. Data validations can be improved if the local developmental and data sets are distributed. Recent advancements of technology, such as Docker allow for local software environments to be preserved using a virtual operating system. Docker allows users to build layers of read/write access files creating a portable operating system which controls exhaustively software versions, data, and systematically preserves one’s complete software environment. Conserving a researcher’s developmental environment advances analytical reproducibility if the workflow is publicly distributed. We suggest a global distributive practice for scholarly publications that regularly includes the virtualized operating system containing all raw analytical data, derived results, and computational software. Currently Docker, compiled sfotware, usually through CMake, and virtual machines are being utilized showing a trend moving toward a global distributive practice linking written methodologies, and supplemental data, to the utilized computational environment.

Comparing Docker as a distributive practice to virtual machines seems roughly equivalent. Distributed virtual machines are easy to download and the environment allows for re-generating resultant calculations. However, this is limited if the research community advances the basic requirements for written methodologies and begins to adopt a large scale virtualized distribution converging to an archive of method environments which would make hosting complete virtual machines impractical or impossible. Whereas archiving Docker containers merely requires a written file as a few bytes in size. If an archive were constructed as a Methods-Analytical-Omnibus, where each research article would link to a distributed methods enviroment, an archive of virtual machines for the entire research community is impossible. However, an archive of Dockerfiles, each containing a few bytes, is quite realistic.

Novel bioinformatic software is often distributed as cross-platform flexible build process independent of compiler which reaches Apple, Windows and Linux users. The scope of novel analytical code is not to manage, nor preserve, computational environments but to have environment independent source code as transportable executables. Docker however does manage operating systems, and the scope for research best practices does include gathering sets of source executables into a single collection, of minimum space, and maximum flexibility that virtual machines can not compete; Docker can provide the ability for the research community to simultaneously advance publication requirements and develop the future computational frameworks in cloud.

Another gained advantage for using Docker as the machine manifesting the practice of reproducible research methods is due to the trend of well-branded organizations such as Illumina’s BaseSpace platform, Google Genomics, or SevenBridges, which all offer bioinformatic computational software structures using Docker as the principle framework. Cloud computational environments offer many advantages over local high-performance in-house computer clusters which systematically structure reproducible methodologies and democratizes medical science. Cloud computational ecosystems preserve an entire developmental environment using the Docker infrastructure improving bioinformatic validation. Containerized cloud applications are instances of the global distributive effort and are favorable compared to local in-house computational pipelines because they offer rapid access to numerous public workflows, easy interfacing to archived read databases, and accelerate the upholding process of raw data. The Google Genomics Cloud has begun to make first steps with integrating cloud infrastructure with the Broad Institute, whereas Illumina’s BaseSpace platform has been hosting novel computational applications since its launch. Google Genomics recently announced a collaboration with the Broad’s software, Genome Analysis Tool Kit (GATK) and now hosts GATK (https://cloud.google.com/genomics/gatk) containerized using Docker. The Docker infrastructure is also utilized by Seven Bridges and Illumina’s BaseSpace Platform and each application is containerized by Docker. For researchers uninterested in designing an exhaustive cloud application, methodology writings can instead publicize containerized workflows with much less effort by providing the Dockerfile which containerizes the corresponding methodology. Scholarly publications that choose only a written method section passively make validation gestures, which is arguably inadequate in comparison to the rising trend or well-branded organizations. We envision a future where published works will share conserved analytic environments, instantiated cloud software accessed by web-distributed methodologies, and/or large databases organizing multitudes of Dockerfiles, with accession numbers, thus strengthening links between raw sequencing data, analytical results, and utilized software.

Cloud computational software does not only wish to crystallize research methods into a pristine pool of transparent methodologies, the other objective is to match the rate of production of high quality analytical results to the rate of production of public data, which reaches hundreds of Petabytes annually. Dr. Atul Butte also observed that with endless public data, the traditional method for practicing science has inverted; no longer does a scientist formulate a question and then experimentally measure observations which test the hypothesis. In the modern area, empirical observations are being made an an unbounded rate, the challenge now is formulating the proper question. Given a near-infinite amount of observations, what is the phenomena that is being revealed? Cloud computational software can accelerate the production of hypotheses by increasing the flexibility of scientific exploration from its efficiency gained by the removal of file transferring and formatting bottle-necks.

Many bioinformaticians have noted a rising trend in biotechnology predicting that open data, and open cloud centers, will democratize research efforts into a more inclusive practice. With the presence of cloud interfacing applications such as Illumina’s BaseSpace Command Line Interface, DNA-Nexus, SevenBridges, and Google Genomics, becoming more popular, cloud environments pioneer the effort for achieving standardized bioinformatic protocols and democratizing research efforts.

Democraticization of big-data efforts has some possible negative consequences. Accessing, networking, and integrating software applications for distributing data as a public effort requires massive amounts of specialized technicians to maintain and develop cloud centers that many research institutions are migrating toward. Currently, it is fairly common for research centers to employ highperformance computer clusters which store laboratory software and data locally; cloud computing clusters are beginning to offer clear advantages compared with local “closed” computer clusters. Collaborations are becoming a more common practice for large research efforts, and sequencing databases have been distributing data globally, making cloud storage more efficient. This implies that services from cloud centers will most likely be offered by a very few elite organizations because the large scale of cloud services will prevent incentives for smaller companies and Moore’s Law will shorten profits of newer technologies. Chris Anderson’s text “The Long Tail” proves that modern economic growth is controlled not by supply, but by the consumer demand as a function of the offerings from “gateway portals” which control accessibility for consumption, moderating and directing consumers toward alternative selections [37]. With regard to recent changes relating to media consumption and e-commerce, democratization allows independent alternative selections more exposure equalizing profits for lower ranked selections “at the tail”. However, it may be possible that the abundant amount of data distributed over storage archives, which stimulates an economically abundant environment, could shift into a fiercely controlled economic environment of scarcity. For bioinformatics, it is very likely that few elite organizations will provide services to cloud computing environments acting as a gateway which directs the global research community toward a narrow set of well-established, standardized, computational applications; thus if a “gold-standard” is reached for computational applications the range of alternative selections could remain non-existent which could diminish the future of bioinformaticians directing them to small garages instead of the technocratic places such as the Silicon Valley motivated not be from a spirit of entrepreneurialism but from a lack of funding. If this holds true, then this implies that the field of applied biostatistics could become completely automated which in turn would reduce the need for analysts polarizing applied research into two pure domains: pure biology and pure mathematics. For instance, limma based differential expression analysis is fully automated over the Gene-Expression-Omnibus’ website where all archived reads can be analyzed using limma and GEO2R software and can be piped into Advaita’s fully automated pathway analysis guide. Automative downstream analyses is not without its drawbacks; most computational software is highly specialized for niche groups with a mathematical framework constructed by specialized assumptions, this would enforce a need for the existence of diverse array of computational developments, and thus a large community of developers. Although the automation of analytical results seems unavoidable, the benefits seem to outweigh the negative consequences.

## 6 Conclusion

Arkas integrates the Kallisto [1] pseudoalingment algorithm into the BioConductor ecosystem that can implement large scale parallel ultra-fast transcript abundance quantification over the BaseSpace Platform. We reduce a computational bottleneck freeing inefficiencies from utilizing ultra-fast transcript abundance calculations while simultaneously connecting an accelerated quantification software to the Sequencing Read Archive. Thus we remove a second bottleneck occurred by reducing the necessity of database downloading; instead we encourage users to download aggregated analysis results. We also expand the range of common sequencing protocols to include an improved gene-set enrichment algorithm, Qusage [19] and allow for exporting into an exhaustive pathway analysis platform, Advaita, over the AWS field in parallel. We encapsulate building annotations libraries for arbitrary fasta files using custom software TxDbLite [4] which may annotate coding, non-coding RNA, with a self generated repeatome that includes genomic repetitive elements such as ALUs, SINEs and/or retro-transposons.

## Acknowledgment

This project was funded by grants from Illumina, Leukemia Lymphoma Society-Quest for Cures 0863-15, and Tower Cancer Research Foundation.

